# Polyamine-controlled proliferation and protein biosynthesis are independent determinants of hair follicle stem cell fate

**DOI:** 10.1101/2020.04.30.070201

**Authors:** Kira Allmeroth, Christine S. Kim, Andrea Annibal, Andromachi Pouikli, Carlos Andrés Chacón-Martínez, Christian Latza, Adam Antebi, Peter Tessarz, Sara A. Wickström, Martin S. Denzel

**Affiliations:** Max Planck Institute for Biology of Ageing, Joseph-Stelzmann-Str. 9b, D-50931 Cologne, Germany; CECAD - Cluster of Excellence, University of Cologne, Joseph-Stelzmann-Str. 26, D-50931 Cologne, Germany; Center for Molecular Medicine Cologne (CMMC), University of Cologne, Robert-Koch-Str. 21, D-50931 Cologne, Germany; Helsinki Institute for Life Science, Biomedicum Helsinki, Haartmaninkatu 8, FI-00290 Helsinki, Finland; Wihuri Research Institute, Biomedicum Helsinki, Haartmaninkatu 8, FI-00290 Helsinki, Finland; Stem Cells and Metabolism Research Program, Faculty of Medicine, Yliopistonkatu 3, FI-00014 University of Helsinki, Finland

**Keywords:** Hair follicle stem cells, mRNA translation, polyamines, cell fate

## Abstract

Stem cell differentiation is accompanied by an increase in mRNA translation. The rate of protein biosynthesis is influenced by the polyamines putrescine, spermidine, and spermine that are essential for cell growth and stem cell maintenance. However, the role of polyamines as endogenous effectors of stem cell fate and whether they act through translational control remains obscure. Here, we investigated the function of polyamines in stem cell fate decisions using hair follicle stem cell (HFSC) organoids. HFSCs showed lower translation rates than progenitor cells, and a forced suppression of translation by direct targeting of the ribosome or through specific depletion of natural polyamines elevated stemness. In addition, we identified N1-acetylspermidine as a novel parallel regulator of cell fate decisions, increasing proliferation without reducing translation. Overall, this study delineates the diverse routes of polyamine metabolism-mediated regulation of stem cell fate decisions.

**Key Points:**

- Low mRNA translation rates characterize hair follicle stem cell (HFSC) state
- Depletion of natural polyamines enriches HFSCs via reduced translation
- N1-acetylspermidine promotes HFSC state without reducing translation
- N1-acetylspermidine expands the stem cell pool through elevated proliferation

## Introduction

Translation of messenger RNAs (mRNAs) is one of the most complex and energy-consuming processes in the cell (Roux & Topisirovic, 2018). It comprises initiation, elongation, termination, and ribosome recycling and plays a key role in gene expression regulation (Hershey et al., 2019). The rate of protein synthesis is tightly controlled by signaling pathways that sense various internal and external stimuli (Roux & Topisirovic, 2018). Deregulation of translation can manifest in a variety of diseases, including neurodegeneration and cancer (Tahmasebi et al., 2018b). The regulation of mRNA translation has also been implicated in early cell fate transitions (Ingolia et al., 2011, Kristensen et al., 2013, Lu et al., 2009). Several studies in both embryonic and somatic stem cells demonstrated that global translation is suppressed in stem cells to retain an undifferentiated state and increased in progenitor cells (Sampath et al., 2008, Signer et al., 2014, Zismanov et al., 2016).

Hair follicle stem cells (HFSCs) represent an excellent paradigm to study adult somatic stem cell fate decisions, since they fuel cyclical rounds of hair follicle regeneration during the natural hair cycle (Blanpain & Fuchs, 2009, Fuchs et al., 2001). Interestingly, Blanco et al. (2016) showed that the activation of HFSCs during the transition from the resting phase, telogen, to proliferation in anagen during the hair cycle also coincides with increased translation. These data demonstrate a functional role for increased translation during differentiation in the epidermis. However, the mechanisms that regulate translation in stem cells in general and upon differentiation remain poorly understood.

One important determinant of translation rates is the availability of the natural polyamines putrescine, spermidine, and spermine. These polycations are essential for cell growth and can bind to a variety of negatively charged cellular molecules, including nucleic acids (Dever & Ivanov, 2018). Polyamine homeostasis is critical to cell survival and is achieved by coordinated regulation at different levels of their synthesis, degradation, uptake and excretion (Wallace et al., 2003). Interestingly, up to 15% of polyamines are stably associated with ribosomes in *Escherichia coli*, highlighting their importance in protein synthesis (Cohen & Lichtenstein, 1960). Additionally, the polyamine spermidine is converted to the amino acid hypusine, which post-translationally modifies the elongation factor eIF5A (Park et al., 1981). This modification is critical for translation elongation, especially for difficult substrates like polyproline (Gutierrez et al., 2013, Saini et al., 2009).

Given that increased polyamine levels positively regulate translation and that high translation rates promote stem cell differentiation, it is surprising that previous studies have implicated elevated polyamine levels in stem cell maintenance. For example, polyamines positively influence expression of MINDY1, a deubiquitinating enzyme, which promotes embryonic stem cell (ESC) self-renewal (James et al., 2018). Additionally, forced overexpression of the rate-limiting enzymes in polyamine biosynthesis adenosylmethionine decarboxylase (AMD1) or ornithine decarboxylase (ODC) in ESCs results in delayed differentiation upon removal of LIF and improves cellular reprogramming (Zhang et al., 2012, Zhao et al., 2012). In line with these observations, differentiation of human bone marrow-derived mesenchymal stem cells coincides with decreased polyamine levels (Tsai et al., 2015). Overall, these data suggest that changes in polyamine levels can affect cell fate. However, the link between polyamines and translation in cell fate decisions remains unclear. Do low polyamine levels endogenously reduce translation rates in stem cells? Does reduction of polyamine levels, and translation, affect stem cell maintenance and function? Might distinct polyamine species differ in their effects on stem cells through translation or other mechanisms?

To address these questions, we used an *in vitro* organoid HFSC culture system and investigated polyamines as key regulators of mRNA translation in stem cell fate decisions. We confirm reduced translation as a key property of HFSCs and demonstrate that forced reduction of translation promotes the stem cell state. We find that HFSCs display low levels of natural polyamines, which implicates polyamines as endogenous gatekeepers of translation in stem cells. Accordingly, specific depletion of natural polyamines using the polyamine analogue N1,N11-diethylnorspermine (DENSpm) improved stem cell maintenance through reduced translation. Surprisingly, depletion of all polyamines by difluoromethylornithine (DFMO) does not affect cell fate. Therefore, we hypothesized that the polyamine pathway can also modulate cell fate independently of reduced translation. Intriguingly, we identify a new function for N1-acteylspermidine (N1-AcSpd) in promoting proliferation, expanding the stem cell pool. Taken together, our data demonstrate that polyamines play a central role in stem cell fate decisions by controlling the key processes translation and proliferation to ensure stem cell maintenance and tissue homeostasis.

## Results

### Low translation rates mark the HFSC state and decreasing translation enhances stemness in the 3D-3C organoids

It has been previously described that stem cells display lower translation rates than their differentiated counterparts (Tahmasebi et al., 2018a). To test if this was also true in epidermal stem cells, we sorted freshly isolated mouse epidermal cells using three different markers (Fig. 1A): α6 integrin is expressed in all progenitors both in the interfollicular epidermis (IFE) and the hair follicle (HF) (Li et al., 1998, Sonnenberg et al., 1991); the hematopoietic stem cell marker CD34 marks the bulge stem cells of the HF (Trempus et al., 2003); stem cell antigen-1 (Sca1) expression can be detected in the infundibulum (IFD) region of the HF and the basal layer of the epidermis (Jensen et al., 2008). By gating for Sca1 negative cells, IFE and IFD progenitors can be separated from HF progenitors. Thus, we compared HF bulge stem cells (Sca1-/α6+/CD34+) to HF outer root sheath cells (Sca1-/α6+/CD34-). We performed puromycin incorporation to quantify the amount of newly synthesized proteins (Schmidt et al., 2009). Western blot analysis revealed a significant reduction of puromycin incorporation in Sca1-/α6+/CD34+ stem cells (Fig. 1B-C), confirming reduced translation rates in HFSCs *in vivo*.

**Figure 1:**
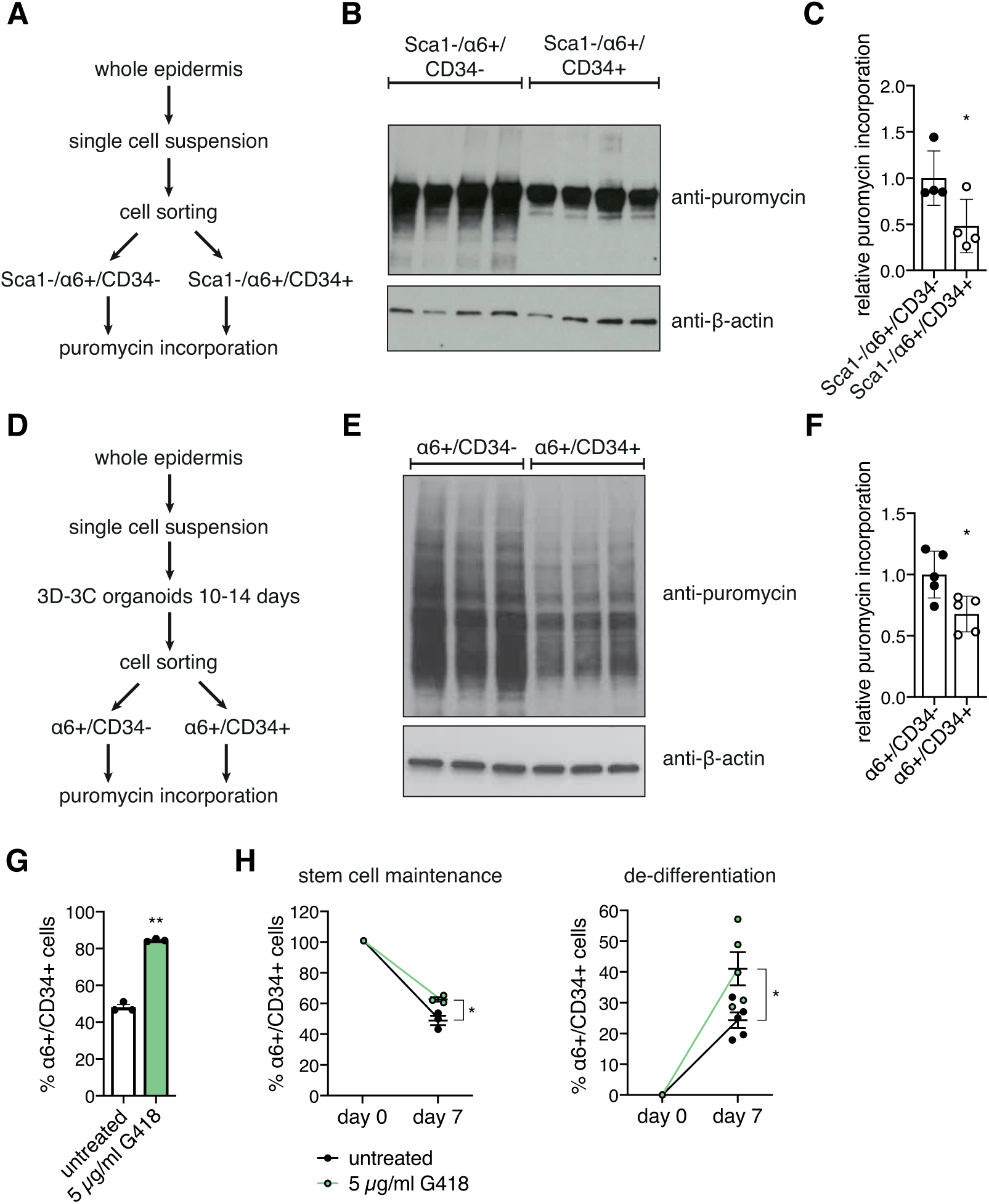
Low translation rates mark the HFSC state and decreasing translation enhances stemness in the 3D-3C organoids. (A) Schematic representation of the workflow for puromycin incorporation using freshly isolated cells. (B) Western blot analysis after puromycin incorporation in Sca1-/α6+/CD34- progenitor cells compared to Sca1-/α6+/CD34+ hair follicle stem cells (HFSCs). (C) Quantification of the Western blot in (B). Mean ± SD (n=4); * p<0.05 (t test). (D) Schematic representation of the workflow for puromycin incorporation using 3D-3C cultured cells. (E) Representative Western blot analysis after puromycin incorporation in α6+/CD34- progenitor cells compared to α6+/CD34+ HFSCs. (F) Quantification of the Western blot in (E). Mean ± SD (n=5); p<0.05 (t test). (G) Ratio of α6+/CD34+ cells after two weeks of 3D-3C culture with or without G418 treatment. Mean ± SEM (n=3); ** p<0.01 (t test). (H) Ratio of α6+/CD34+ cells at day 0 and day 7 post-sorting starting from 100 % α6+/CD34+ cells (left) or 100 % α6+/CD34- cells (right) with or without G418 treatment. Mean ± SEM (n≥3); * p<0.05 (t test).

To understand the mechanisms and functional consequences of this difference in translation, we made use of an *ex vivo* organoid culture system (3D-3C culture) established by Chacon-Martinez et al. (2017) allowing for the long-term culture and manipulation of HFSCs and their direct progeny. Importantly, in this culture cells maintain self-renewal capacity and multipotency and respond to the same signals as they do *in vivo*. In the 3D-3C organoids, a balance between α6+/CD34+ HFSCs and α6+/CD34- progenitor cells is formed in a self-driven process. Based on their transcriptome and marker expression analysis, these progenitor cells represent HF outer root sheath (ORS) cells and inner bulge cells (Kim et al., 2019), both of which represent HFSC progeny and act as niche cells for HFSCs *in vivo* (Hsu et al., 2014). To confirm cell identity of α6+/CD34+ HFSCs and α6+/CD34- ORS progenitors, we sorted cells after two weeks of 3D-3C culture and analyzed gene expression by RNA-sequencing. Comparing the two populations, the most enriched GO term was skin development (Fig. S1A). As expected, the differentiation markers Krt1 and Krt10 were enriched in progenitor cells, while the expression of the stem cell marker genes Id2, Lhx2, and Sox9 was higher in HFSCs (Fig. S1B). Thus, α6+/CD34- progenitors and α6+/CD34+ HFSCs are clearly distinguishable at the gene expression level and represent distinct cellular states. Next, we investigated puromycin incorporation in the two cell populations after sorting and found reduced translation in α6+/CD34+ HFSCs (Fig. 1D-F). Thus, the 3D-3C organoids maintain this key *in vivo* property of HFSCs, making them a suitable model to study the influence of translation on cell fate.

Next, we asked whether reduced translation might drive cell fate decisions in HFSCs. To this end, we manipulated translation in the 3D-3C organoids by addition of low doses of G418, a well described inhibitor of translation elongation (Bar-Nun et al., 1983). We confirmed reduced translation after 4 h of G418 treatment in the 3D-3C organoids by measuring puromycin incorporation (Fig. S1C-D). Strikingly, a forced decrease in translation resulted in a significant increase of α6+/CD34+ stem cells after two weeks of 3D-3C culture (Fig. 1G). To understand whether this increase had a functional consequence, we analyzed stem cell potency in a colony formation assay. For this, cells were seeded in clonal density on a feeder layer and cultured for 2 to 3 weeks (Fig. S1E). Quantification of total colony number revealed a significant increase upon G418 treatment in the organoid culture, indicating that inhibiting translation increased stem cell proliferative potential (Fig. S1F-G). The self-organizing plasticity of the organoids results in a balance between α6+/CD34- and α6+/CD34+ cells (Chacon-Martinez et al., 2017). This balance is influenced by self-renewal and differentiation of HFSCs but also by proliferation and de-differentiation of progenitors back to the stem cell state. Thus, to investigate if the increase in stem cell potency was accompanied by increased self-renewal and de-differentiation in the G418-treated organoids, we sorted the cells after two weeks of 3D-3C culture to generate pure populations of either α6+/CD34- or α6+/CD34+ cells. These pure populations were cultured for one week with or without G418 treatment before analysis. Interestingly, G418 treatment prevented stem cell differentiation and promoted de-differentiation of progenitors back to the stem cell state (Fig. 1H). Taken together, these data suggest that reduced translation is not merely a consequence of stemness. Instead, decreasing translation actively promotes the stem cell state.

### Depletion of natural polyamines is a feature of stem cells and can drive stemness *in vitro*

To further analyze the effect of decreased translation rates on cell fate, we focused on the polyamine pathway (Fig. 2A). Reduction of the natural polyamines putrescine, spermidine, and spermine has been implicated in reduced translation (Dever & Ivanov, 2018, Pegg, 2016). We thus asked whether stem cells might display lower polyamine levels compared to their differentiated counterparts. Indeed, when we sorted cells after two weeks of 3D-3C culture and measured polyamines by LC-MS, we found reduced levels of putrescine, spermidine, and spermine in α6+/CD34+ HFSCs compared to α6+/CD34- progenitors (Fig. 2B).

**Figure 2:**
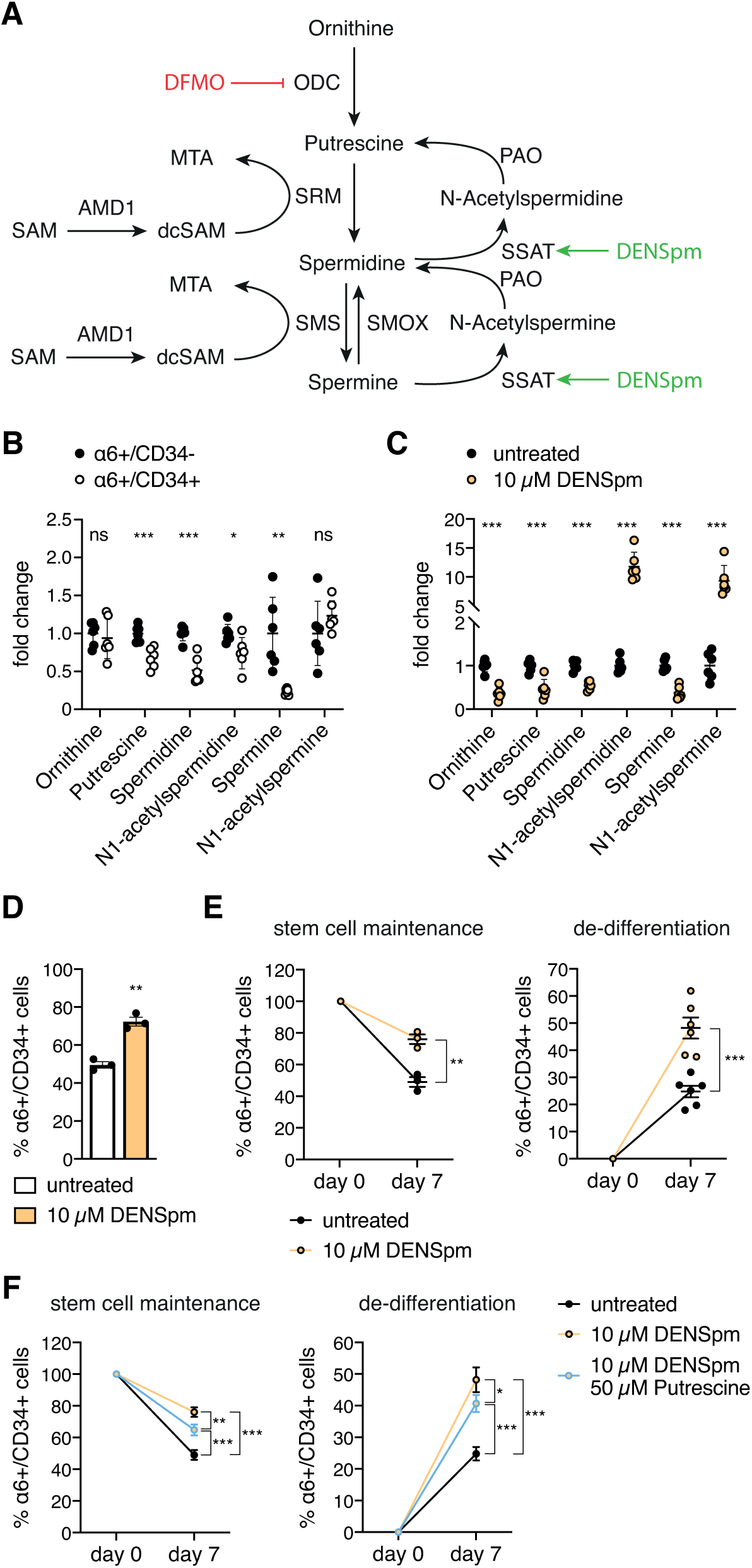
Reduction of translation by depletion of natural polyamines recapitulates the stem cell state and regulates cell fate decisions. (A) Schematic representation of the polyamine pathway. DENSpm is marked in green and DFMO is depicted in red. (B) Polyamine levels in sorted α6+/CD34- and α6+/CD34+ cells after 3D-3C culture. Mean ± SEM (n=6). (C) Polyamine levels in 3D-3C cultured cells with and without DENSpm treatment. Mean ± SEM (n=6). (D) Ratio of α6+/CD34+ cells after two weeks of 3D-3C culture with or without DENSpm treatment. Mean ± SEM (n=3). (E) Ratio of α6+/CD34+ cells at day 0 and day 7 post-sorting starting from 100 % α6+/CD34+ cells (left) or 100 % α6+/CD34- cells (right) with or without DENSpm treatment. Mean ± SEM (n≥3). (B-E) Statistical significance was calculated by t test. *** p<0.001, ** p<0.01, * p<0.05, ns: not significant. (F) Ratio of α6+/CD34+ cells at day 0 and day 7 post-sorting starting from 100 % α6+/CD34+ cells (left) or 100 % α6+/CD34- cells (right) with or without DENSpm or additional putrescine treatment. Mean ± SEM (n≥3). Two-way ANOVA Dunnett post-test. *** p<0.001, ** p<0.01, * p<0.05.

To probe polyamine function in stem cell fate decisions, we manipulated their levels using the spermine analogue N1,N11-diethylnorspermine (DENSpm), which increases spermidine/spermine N1-acetyltransferase (SSAT) expression (Coleman et al., 1995, Fogel-Petrovic et al., 1996, Parry et al., 1995). During polyamine catabolism, SSAT acetylates spermidine and spermine using acetyl-CoA to form N1-acetylspermidine and N1-acetylspermine, respectively (Fig. 2A). Thus, DENSpm depletes natural polyamines, while the acetylated forms accumulate (Uimari et al., 2009), resulting in decreased mRNA translation (Landau et al., 2010). LC-MS analysis confirmed reduced natural polyamines and elevated acetylated forms upon DENSpm treatment in the 3D-3C organoids (Fig. 2C). Strikingly, supplementation with DENSpm resulted in a significant increase in α6+/CD34+ stem cells after two weeks of 3D-3C culture (Fig. 2D). The increase in stemness was confirmed in the colony formation assay that revealed an elevated colony number upon DENSpm treatment in the 3D-3C organoids, suggesting increased proliferative potential (Fig. S2A-B). To investigate stem cell maintenance and de-differentiation separately, pure populations were sorted and cultured for one week with or without DENSpm treatment. Both mechanisms were affected by the depletion of natural polyamines, increasing α6+/CD34+ stem cells (Fig. 2E). To rule out that the changes in stem cell number and proliferative potential were secondary to an effect on cell proliferation, we measured EdU incorporation in both α6+/CD34+ and α6+/CD34- cells. EdU incorporation was not affected by DENSpm treatment in either of the cell populations (Fig. S2C). Additionally, we analyzed apoptosis by annexin V staining. While annexin V staining was not changed in α6+/CD34+ cells, DENSpm treatment slightly increased apoptosis in α6+/CD34- progenitors (Fig. S2D). However, this effect was not strong enough to account for the increase in α6+/CD34+ HFSCs in treated organoid cultures. Furthermore, we demonstrated increased stem cell potency in the colony-forming assay (Fig. S2A-B). Thus, our data indicate that the promotion of the stem cell state by DENSpm was through a direct effect on stem cell fate. Since the effect of DENSpm supplementation on the pure stem cell population was comparable to G418 treatment (Fig. 1H), we speculated that reduced translation might be the underlying mechanism of increased stemness. To test this hypothesis, we supplemented DENSpm treated cells with putrescine to elevate the natural polyamines. Indeed, this double treatment showed a partial rescue as stem cell maintenance and de-differentiation were significantly suppressed (Fig. 2F). Taken together, these data demonstrate that α6+/CD34+ stem cells have lower polyamine levels, and that the reduction of natural polyamines influences HFSC fate through decreased translation.

### N1-acetylspermidine is a novel determinant of HFSC fate acting independently of reduced translation

To further test if reduction of translation by depletion of natural polyamines caused the observed cell fate changes upon DENSpm treatment, we made use of difluoromethylornithine (DFMO), an irreversible ODC inhibitor (Metcalf et al., 1978) (Fig. 2A) that depletes all polyamines. In contrast to ODC inhibition by DFMO, DENSpm supplementation caused specific reduction of the natural polyamines putrescine, spermidine, and spermine, while the acetylated forms accumulated (Fig. 2C). Surprisingly, DFMO treatment did not affect cell fate in the 3D-3C organoids (Fig. 3A). Thus, we hypothesized that reduced mRNA translation through decreased polyamine availability was not the only determinant of the cell fate changes observed upon DENSpm treatment. Since additional putrescine supplementation only partially rescued the DENSpm effect on stem cell maintenance and de-differentiation, we speculated that putrescine itself might affect cell fate. Indeed, putrescine treatment in the 3D-3C organoids resulted in an increase in the number of α6+/CD34+ stem cells (Fig. 3B). Increased stemness was confirmed by elevated colony-forming ability, showing enhanced proliferative potential (Fig. S3A-B). Surprisingly, putrescine supplementation specifically enhanced de-differentiation of α6+/CD34- progenitors to α6+/CD34+ HFSCs, while stem cell maintenance was not affected (Fig. 3C). However, the effect was not as pronounced as with DENSpm treatment (Fig. 2E). To further investigate the effect of putrescine supplementation, we measured intracellular polyamine levels in the 3D-3C organoids. Spermidine and N1-acetylspermidine (N1-AcSpd) levels were increased in response to putrescine treatment (Fig. 3D). In addition, an elevation of putrescine confirmed the effectiveness of our treatment. Since both DENSpm and putrescine supplementation led to elevated N1-AcSpd levels (Fig. 2C and Fig. 3D), we speculated that increased N1-AcSpd might cause the cell fate changes. An increase in any polyamine would not be expected to reduce translation and would then be mediated through a distinct mechanism. Intriguingly, N1-AcSpd treatment significantly increased the number of α6+/CD34+ stem cells in the 3D-3C organoids (Fig. 3E). Stemness was confirmed by enhanced proliferative potential in the colony formation assay (Fig. S3C-D). As expected, analysis of puromycin incorporation confirmed that the effect on stemness occurred without a reduction of translation. Instead, translation rates were increased upon short-term N1-AcSpd treatment (72 h) (Fig. S3E-F), which did not affect the ratio of HFSCs in the 3D-3C organoids (Fig. S3G). Next, we measured intracellular polyamine levels upon N1-AcSpd treatment and found that apart from N1-AcSpd, also putrescine was increased (Fig. 3F). Thus, the effect of N1-AcSpd treatment on intracellular polyamines resembled the putrescine supplementation (Fig. 3D). However, in contrast to putrescine addition, N1-AcSpd treatment enhanced not only de-differentiation, but also stem cell maintenance when starting from pure populations (Fig. 3G). In stark contrast, supplementation with N1-acetylspermine (N1-AcSpm) did not influence the ratio of HFSCs (Fig. 3H). Thus, elevation of polyamines in general was not sufficient; rather, putrescine and N1-AcSpd treatment had specific effects on cell fate. Overall, these data implicate N1-AcSpd as a novel cell fate regulator that acts independently of reduced translation.

**Figure 3:**
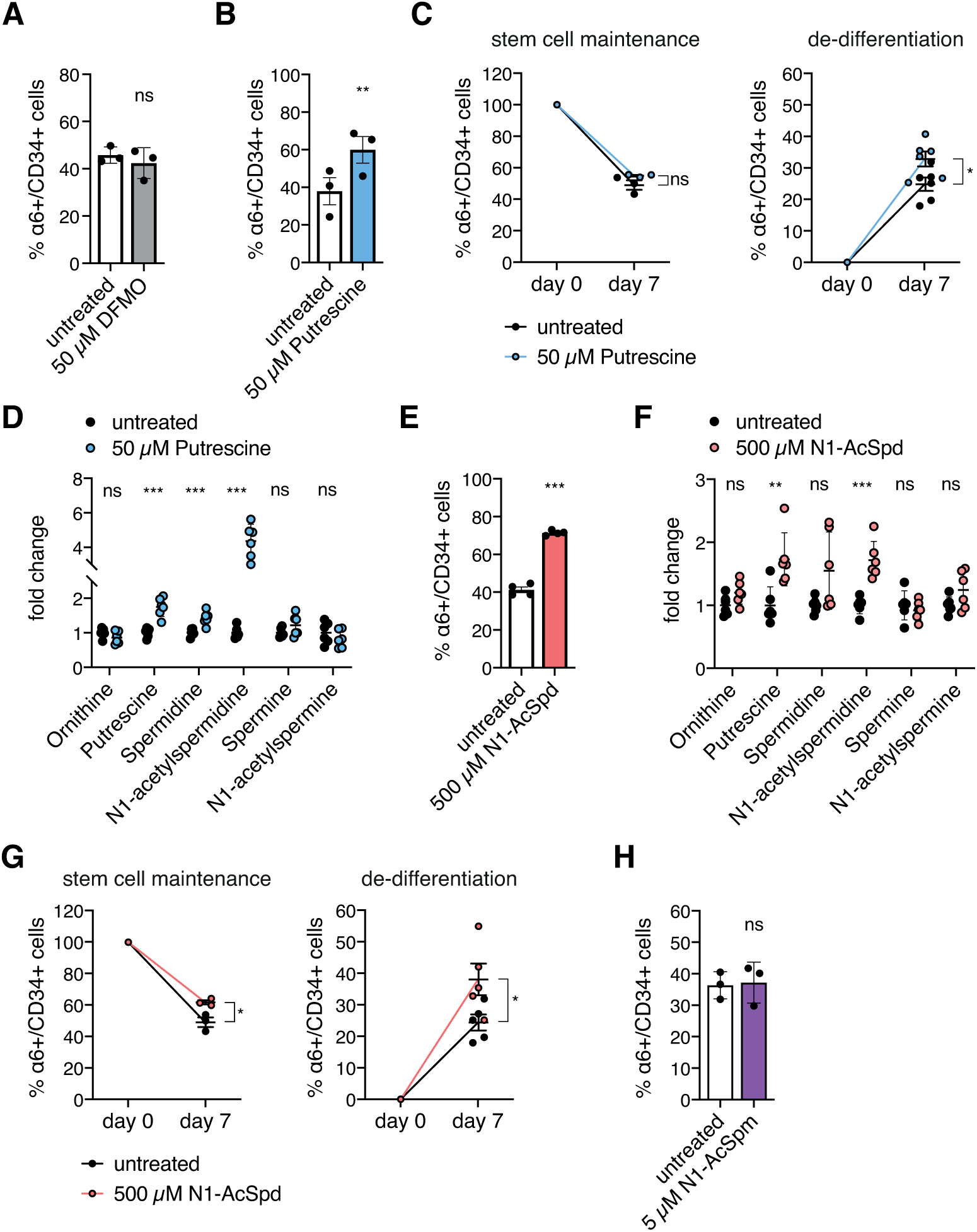
N1-acetylspermidine is a novel regulator of hair follicle stem cell fate. (A) Ratio of α6+/CD34+ cells after two weeks of 3D-3C culture with or without DFMO treatment. Mean ± SEM (n=3). (B) Ratio of α6+/CD34+ cells after two weeks of 3D-3C culture with or without putrescine treatment. Mean ± SEM (n=3). (C) Ratio of α6+/CD34+ cells at day 0 and day 7 post-sorting starting from 100 % α6+/CD34+ cells (left) or 100 % α6+/CD34- cells (right) with or without putrescine treatment. Mean ± SEM (n≥3). (D) Polyamine levels in 3D-3C cultured cells with and without putrescine treatment. Mean ± SEM (n=6). (E) Ratio of α6+/CD34+ cells after two weeks of 3D-3C culture with or without N1-AcSpd treatment. Mean ± SEM (n=3). (F) Polyamine levels in 3D-3C cultured cells with and without N1-AcSpd treatment. Mean ± SEM (n=6). (G) Ratio of α6+/CD34+ cells at day 0 and day 7 post-sorting starting from 100 % α6+/CD34+ cells (left) or 100 % α6+/CD34- cells (right) with or without N1-AcSpd treatment. Mean ± SEM (n≥3). (H) Ratio of α6+/CD34+ cells after two weeks of 3D-3C culture with or without N1-AcSpm treatment. Mean ± SEM (n=3). ns: not significant (t test).

### N1-acetylspermidine treatment affects hair follicle stem cell fate by increasing proliferation

To investigate the molecular mechanism by which N1-AcSpd supplementation promotes stemness, we performed RNA-sequencing of 3D-3C cultured cells upon short-term N1-AcSpd treatment (72 h). HFSCs and progenitor cells were purified and analyzed separately (Fig. S4A). We compared α6+/CD34- cells with α6+/CD34+ cells to identify differentially expressed genes for control (CTR) and N1-AcSpd treatment (Fig. S4B). The two groups of differentially expressed genes were subjected to further analysis. Around 2.200 genes with differential expression between cell types were not affected by N1-AcSpd treatment (Fig. S4C). 851 genes were differentially expressed between HFSCs and progenitors only in the presence of N1-AcSpd, while 540 genes were differentially expressed between cell types exclusively in untreated control cells (Fig. S4C). GO term analysis was performed with the three different groups using Metascape (Zhou et al., 2019). Strikingly analysis of the genes differentially expressed between HFSCs and progenitors only during N1-AcSpd treatment revealed that most of the ten highest ranked GO terms were linked to cell cycle progression (Fig. 4A). The most significant GO term was cell division, of which 54 genes were present among the 851 differentially expressed genes in N1-AcSpd treatment (Fig. S4D). Interestingly, expression of the majority of these 54 genes was higher in α6+/CD34+ stem cells (Fig. 4B). Importantly, N1-AcSpd treatment affected the amplitude of the fold change; the expression of these genes was higher also in the untreated α6+/CD34+ stem cells (Fig. 4B). Consistently, EdU incorporation was higher in untreated α6+/CD34+ HFSCs compared to progenitor cells (Fig. S4E). Together, these data suggest that N1-AcSpd selectively promoted self-renewal of HFSCs. GO term analysis of the around 2.200 genes that differed between HFSCs and progenitors without being affected by N1-AcSpd revealed skin development as the most significant GO term (Fig. S4F). This suggests that N1-AcSpd treatment did not affect general cell identity in the 3D-3C organoids. Consistently, a comparison of the gene expression changes between cell types in a heat map showed a clear separation of α6+/CD34+ HFSCs and α6+/CD34- progenitors (Fig. S4G). Although N1-AcSpd treated cells clustered separately of control cells, their gene expression was very similar to the respective cell type in the control (Fig. S4G). Analysis of the 540 genes differentially expressed between cell types only in untreated cells showed low significance for all GO terms. The most significant GO terms were intraciliary transport and cilium organization (Fig. S4H).

**Figure 4:**
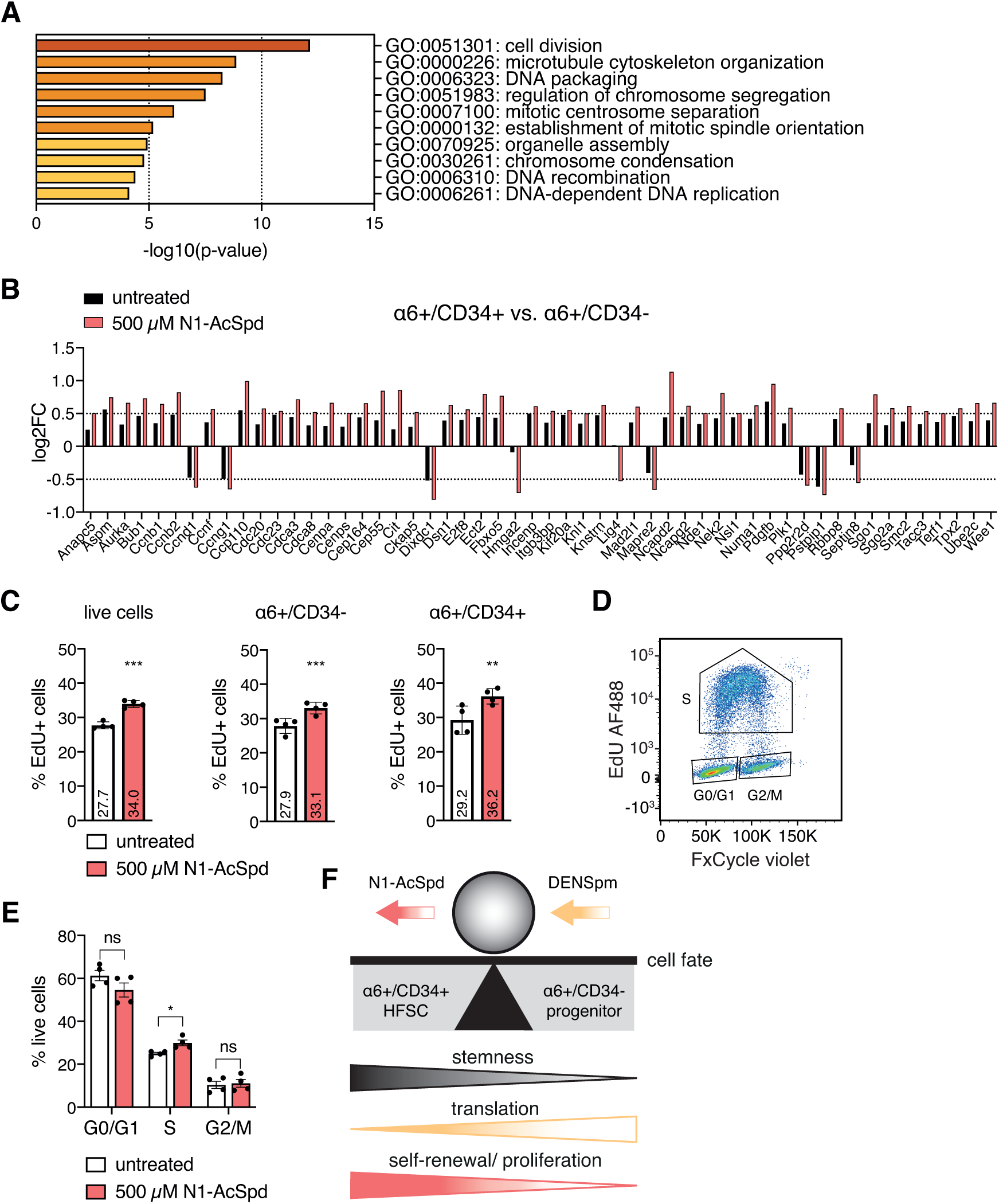
N1-acetylspermidine treatment affects cell fate by increasing cell cyle progression. (A) GO term analysis of differentially regulated genes between cell types (p-value < 0.05, log2FC > ±0.5) only upon N1-AcSpd treatment (depicted in red in Fig. S4A, biological process, metascape.org). (B) log2FC of the genes associated with the GO term “cell division” in untreated and N1-AcSpd treated cells. The ratio of the log2FC in α6+/CD34+ vs. α6+/CD34- cells is shown. (C) Ratio of EdU+ cells with or without N1-AcSpd treatment (72 h). EdU was incorporated for 2 h. Mean ± SEM (n=4). *** p<0.01; ** p<0.01 (t test). (D) Dot plot of FxCycle violet intensity and EdU AF488 intensity, separating the phases of the cell cycle (G0/G1, S, G2/M). (E) Ratio of live cells with or without N1-AcSpd treatment (72 h) according to their cell cycle phase distribution. Mean ± SEM (n=4). * p<0.05; ns: not significant (t test). (F) Model of polyamine-controlled effects on cell fate. Manipulation of translation or proliferation by DENSpm or N1-AcSpd, respectively, results in the enrichment of HFSCs.

Since the analysis of global gene expression changes clearly indicated that N1-AcSpd treatment would affect proliferation, we measured EdU incorporation. Importantly, short-term N1-AcSpd supplementation (72 h) increased the number of EdU+ cells (Fig. 4C). The effect on α6+/CD34+ HFSCs was higher compared to α6+/CD34- progenitor cells (Fig. 4C). Still, consistent with increased de-differentiation and elevated stem cell maintenance, proliferation of both populations was affected by the treatment. Furthermore, we analyzed the cell cycle phase distribution of the cells using FxCycle violet staining. Plotting of FxCycle violet intensity on the x-axis and EdU AF488 intensity on the y-axis enabled clear separation of the different cell cycle phases (Fig. 4D). Consistent with increased EdU incorporation, we observed faster cell cycle progression with fewer cells in G0/G1 phase and more cells in S phase upon short-term N1-AcSpd treatment (72 h) (Fig. 4E). The number of cells in G2/M phase was not changed. Taken together, these data suggest that the acetylated polyamine N1-AcSpd fulfills a functional role in cell cycle progression and that it affects cell fate by increasing proliferation. Overall, in this study we decipher the different and independent polyamine-controlled effects on HFSC fate. We demonstrate that decreasing translation rates by the depletion of natural polyamines through DENSpm, as well as enhancing proliferation by N1-AcSpd shifts the balance towards the stem cell state in the 3D-3C organoids (Fig. 4F).

## Discussion

In this study we delineate the different routes of polyamine-mediated regulation of HFSC maintenance and function. First, we show that reduced translation is not only a marker of the stem cell state. Instead, a forced decrease of translation was sufficient to enhance stemness. Next, we noted low polyamine levels in α6+/CD34+ stem cells and depletion of the natural polyamines with DENSpm resulted in enhanced HFSC maintenance. Strikingly, full depletion of polyamines by DFMO did not recapitulate our findings with DENSpm, suggesting that reduced translation was not the only relevant consequence of polyamine changes in cell fate. Indeed, putrescine supplementation had a beneficial effect on progenitor cell de-differentiation. This was mediated by N1-AcSpd that we identify as novel regulator of HFSC fate, acting independently of reduced translation by increasing proliferation.

Previously, Blanco et al. (2016) showed that translation becomes elevated during the hair cycle growth phase when cells become activated and differentiate. In contrast, quiescent HFSCs in the resting phase show low translation. This agrees with other studies showing that translation is upregulated during differentiation (Sampath et al., 2008, Signer et al., 2014, Zismanov et al., 2016). Importantly, we can reproduce these findings *in vivo* and in the 3D-3C HFSC organoid culture; α6+/CD34+ stem cells in the organoids recapitulate key features of HFSCs *in vivo*. Strikingly, forced inhibition of translation by loss of NOP2/Sun RNA methyltransferase family member 2 (NSUN2) in the mouse blocks the differentiation of HFSCs to progenitor cells (Blanco et al., 2011), clearly showing that upregulation of translation is necessary for differentiation in the HF. Using G418 treatment or the depletion of natural polyamines to reduce translation in the organoids, we confirm promotion of the stem cell state. We further show that the number of stem cells was elevated due to enhanced self-renewal rather than attenuated differentiation using EdU incorporation during DENSpm treatment.

Additionally, we see an effect on progenitor cell de-differentiation, further supporting the idea that a forced decrease of translation is sufficient to promote stemness in the 3D-3C organoids.

Previously, the acetylated polyamines were mostly described as the major group of polyamines exported from the cell (Seiler & Dezeure, 1990). Extending this notion, we find that the intracellular accumulation of N1-AcSpd has an effect on stemness. This occurs without reducing translation, which is consistent with previous observations that acetylated polyamines have no effect on protein synthesis *in vitro* (Kakegawa et al., 1991). Instead, RNA-sequencing of purified cell populations from the organoids revealed that N1-AcSpd treatment affected expression of cell cycle-associated genes. Interestingly, these same genes were found expressed at higher levels in α6+/CD34+ stem cells, which also display higher EdU incorporation compared to progenitor cells in the organoid cultures. Importantly, increased proliferation and stemness have been linked before. A study in ESCs suggests that elevated CDK activity, and thus increased cell cycle progression, contributes to stem cell maintenance (Liu et al., 2017). Strikingly, several studies have described that cells in the G1 phase are more susceptible to cell fate changes and that differentiation is associated with G1 phase lengthening (Calder et al., 2013, Clegg et al., 1987, Sela et al., 2012). Consistently, reprogramming efficiency seems to be linked to successful acceleration of the cell cycle (Guo et al., 2014, Ruiz et al., 2011). Of note, forced overexpression of AMD1 or ODC in mouse fibroblasts, resulting in the accumulation of polyamines, improves reprogramming efficiency (Zhang et al., 2012, Zhao et al., 2012). Our results suggest that this effect is caused by increased proliferation. Collectively, these data indicate that both stem cell maintenance and de-differentiation might be improved by elevated cell cycle progression upon N1-AcSpd treatment in our 3D-3C organoid culture.

Polyamines have been implicated in cell cycle progression before. Under normal growth conditions, polyamine levels are dynamically regulated during the cell cycle (Sunkara et al., 1981), which is due to cyclical changes in activity of ODC and AMD1 (Fredlund et al., 1995). Consistently, several studies have shown that polyamine depletion results in cell cycle arrest (Odenlund et al., 2009, Ray et al., 1999, Yamashita et al., 2013). Intriguingly, ODC activity was shown to be increased by N1-AcSpd, while N1-AcSpm, which did not influence cell fate in the organoid culture, does not affect ODC activity (Canellakis et al., 1989). Thus, activation of ODC might be an important step in cell fate regulation via increased cell cycle progression. Strikingly, expression of ODC is also dynamically regulated during the hair cycle. Nancarrow et al. (1999) showed that ODC is abundantly expressed in proliferating bulb cells of anagen follicles, while entry of the follicle into catagen is accompanied by down-regulation of ODC. These data implicate that polyamine levels, which are controlled by ODC, might track proliferation rates in the hair follicle. Additionally, the authors revealed expression of ODC in ORS cells in vicinity of the follicle bulge region (Nancarrow et al., 1999). After the anagen growth phase, some ORS cells return to the bulge region, where they resume quiescence and CD34 expression. These cells become the primary stem cells for the initiation of the next hair cycle (Hsu et al., 2011). This process is comparable to the de-differentiation of α6+/CD34- progenitor cells back to the stem cell state in the 3D-3C organoids, which was enhanced by putrescine and N1-AcSpd supplementation. Overall, these data suggest that ODC activity, cell cycle progression, and cell fate are tightly linked. In the HF, quiescent stem cells ensure reduced translation rates by low polyamine levels, which increase upon activation due to elevated ODC activity, resulting in accelerated cell cycle progression. Consistently, skin-specific ODC overexpression results in hair loss beginning 2 to 3 weeks after birth, demonstrating the need for low polyamine levels in HFSC quiescence (Soler et al., 1996). Our data suggest that enhanced cell cycle progression promotes stem cell self-renewal upon activation, while differentiation is decreased. At the same time, cell cycle progression might be required for successful de-differentiation of ORS cells back to HFSCs, a notion supported by elevated ODC expression in ORS cells.

Overall, this study dissects independent routes of polyamine-controlled regulation of cell fate: While depletion of natural polyamines endogenously reduced translation, addition of N1-AcSpd accelerated cell cycle progression. Previously, acetylated polyamines were described primarily as the major polyamine species exported from the cell. Here, we demonstrate that N1-AcSpd directly influenced cell fate decisions via increased proliferation. Thus, our study explains why elevated polyamine levels and low translation rates are not mutually exclusive in stem cell maintenance. Instead, they regulate different aspects of cell fate. While low translation rates favor quiescence *in vivo*, enhanced cell cycle progression ensures stem cell self-renewal upon activation. ODC activity might function as the molecular switch to regulate polyamine availability and thus cell fate in the HF. Over the long-term, N1-AcSpd treatment could be a viable intervention to tackle age-associated diseases caused by a decline in tissue homeostasis due to stem cell exhaustion.

## Methods

### Mouse husbandry

Animals were housed on a 12:12 h light:dark cycle with *ad libitum* access to food under pathogen-free conditions in individually ventilated cages. All animals were kept in C57BL/6J background. Animal care and experimental procedures were in accordance with the institutional and governmental guidelines.

### Isolation of epidermal cells

Isolation of epidermal cells from mice in telogen stage was performed as described previously (Chacon-Martinez et al., 2017). In brief, mouse skin was incubated on 0.8 % trypsin (ThermoFisher Scientific) for 50 min at 37 °C. The skin was transferred to 8 ml KGM medium and the epidermis was separated from the dermis. After centrifugation, the keratinocytes were resuspended in ice cold KGM medium and embedded in growth factor-reduced matrigel (Corning Inc.).

### Culture of hair follicle stem cell organoids

Hair follicle stem cell organoids were cultured in KGM medium: MEM medium (Spinner’s modification, Sigma-Aldrich), supplemented with 8 % fetal bovine serum (chelated, ThermoFisher Scientific), penicillin/streptavidin (ThermoFisher Scientific), L-glutamine (ThermoFisher Scientific), insulin (5 µg/ml, Sigma-Aldrich), hydrocortisone (0.36 µg/ml, Calbiochem), EGF (10 ng/ml, Sigma-Aldrich), transferrin (10 µg/ml, Sigma-Aldrich), phosphoethanolamine (10 µM, Sigma-Aldrich), ethanolamine (10 µM, Sigma-Aldrich), CaCl_2_ (14.5 µM, Sigma-Aldrich). 5 µM Y27632, 20 ng/ml mouse recombinant VEGF, 20 ng/ml human recombinant FGF2 (all Miltenyi Biotech) were added to KGM medium. The cells were grown at 37 °C in 5 % CO_2_.

### Cell maintenance

NIH3T3 fibroblasts were grown in DMEM containing 4.5 g/L glucose supplemented with 10 % fetal bovine serum and penicillin/streptavidin (all ThermoFisher Scientific) at 37 °C in 5 % CO_2_ on non-coated tissue culture plates.

### Colony formation assay

NIH3T3 fibroblasts were seeded on collagen G-coated (30 µg/ml in PBS, Biochrom AG) tissue culture plates. The cells were grown at 37 °C in 5 % CO_2_ for two days before proliferation was inhibited with mitomycin C (Sigma-Aldrich).

Hair follicle stem cells were grown in 3D-3C organoids for two weeks and analyzed by flow cytometry. 4000 cells were seeded per 6-well on the fibroblast layer in MEM/HAM’s F12 (FAD) medium with low Ca^2+^ (50 µM, Biochrom AG), supplemented with 10% fetal bovine serum (chelated, ThermoFisher Scientific), penicillin/streptavidin (ThermoFisher Scientific), L-glutamine (ThermoFisher Scientific), ascorbic acid (50 µg/ml, Sigma-Aldrich), adenine (0.18 mM, Sigma-Aldrich), insulin (5 µg/ml, Sigma-Aldrich), hydrocortisone (0.5 µg/ml, Sigma-Aldrich), EGF (10 ng/ml, Sigma-Aldrich), and cholera enterotoxin (10 ng/ml, Sigma-Aldrich). The cells were grown for 2-3 weeks at 32 °C in 5 % CO_2_ until colonies were formed. Remaining fibroblasts were removed using 0.25 % trypsin-ETDA (ThermoFisher Scientific) for 2 min at 37 °C. Trypsin was stopped using supplemented DMEM. After washing the plates with PBS, keratinocytes were fixed with 4 % PFA (Sigma-Aldrich) for 15 min at RT. The cells were washed with PBS twice and the colonies were stained using 1 % crystal violet (Sigma-Aldrich in PBS) for 1 h at RT on an orbital shaker. The wells were washed with tap water until no stain was released and air dried. The plates were scanned and the number of colonies was counted manually.

### Flow cytometry

Matrigel droplets were degraded in 0.5 % trypsin, 0.5 mM EDTA in PBS for 8 min at 37 °C. Trypsin was neutralized using cold KGM. After centrifugation for 5 min at 2000 rpm cells were washed with FACS buffer (2 % FBS, 2 mM EDTA in PBS). Surface marker staining was performed for 30 min on ice with the following antibodies: CD34-eFluor 660 (eBioscience, clone RAM34, 1:100) and ITGA6-PE/Cy7 (Biolegend, clone GoH3, 1:1000) for sorting or CD34-eFluor 660 (eBioscience, clone RAM34, 1:100) and ITGA6-pacific blue (Biolegend, clone GoH3, 1:200) for analysis. Freshly isolated keratinocytes were stained additionally with Sca1-Pacific blue (Biolegend, clone D7, 1:400). 7AAD or PI was used to exclude dead cells. Cells were analyzed on FACSCantoII (BD Biosciences) or sorted on FACSAria IIIu Svea and FACSAria Fusion sorters (BD Biosciences). Sorted cells were collected in 15 ml tubes containing KGM at 4 °C.

### EdU incorporation and detection

Cell proliferation was assessed using the Click-iT™ Plus EdU Alexa Fluor™ 488 Flow Cytometry Assay Kit (ThermoFisher) following the manufacturer’s instructions. Cells were grown for 9 days in 3D-3C culture before 10 µM EdU was added to the medium and incubated for 2 h or 24 h. Matrigel droplets were degraded, the cells were washed with PBS and subsequently stained with fixable LIVE/DEAD-violet (ThermoFisher, 1:500) for 20 min at RT. The cells were washed with FACS buffer and surface marker staining was performed for 30 min on ice using CD34-eFluor 660 (eBioscience, clone RAM34, 1:100) and ITGA6-PE/Cy7 (Biolegend, clone GoH3, 1:1000). The cells were washed with 1 % BSA in PBS, fixed with 4 % PFA for 10 min at RT and permeabilized using Click-iT saponin-based permeabilization buffer for 15 min at RT. The EdU reaction cocktail was prepared following the manufacturer’s instructions and incubated for 30 min at RT in the dark after washing the cells. The cells were analyzed on FACSCantoII.

### Cell cycle analysis

Cell proliferation was assessed using the Click-iT™ Plus EdU Alexa Fluor™ 488 Flow Cytometry Assay Kit (ThermoFisher) as described above. Cells were grown for two weeks in 3D-3C culture before 10 µM EdU was added to the medium and incubated for 2 h. Matrigel droplets were degraded, the cells were washed with PBS and stained with fixable Zombie NIR dye (Biolegend, 1:500) for 20 min at RT. The cells were washed, fixed, and permeabilized and the EdU reaction cocktail was incubated as described above. Subsequently, the cells were permeabilized using 0.1 % triton X-100 in PBS for 15 min at RT and stained with FxCycle violet (ThermoFisher, 1:500) for 30 min at RT. The cells were analyzed on FACSCantoII without further washing.

### Annexin V staining

Matrigel droplets were degraded in 0.5 % trypsin, 0.5 mM EDTA in PBS for 8 min at 37 °C. Trypsin was neutralized using cold KGM. After centrifugation for 5 min at 2000 rpm cells were washed with FACS buffer. Surface marker staining was performed for 30 min on ice using CD34-eFluor 660 (eBioscience, clone RAM34, 1:100) and ITGA6-pacific blue (Biolegend, clone GoH3, 1:200). Cells were washed with PBS and stained with annexin V-AF488 (ThermoFisher Scientific, 1:20 in 100 µl annexin-binding buffer per sample) for 15 min at RT. 100 µl annexin-binding buffer containing PI were added per sample. The cells were analyzed on FACSCantoII.

### Puromycin incorporation

Cells were incubated with 10 µg/ml puromycin in the medium for exactly 10 min at 37 °C. The cells were washed with PBS and collected in RIPA buffer (50 mM TrisHCl, 120 mM NaCl, 0.1 % SDS, 1 % NP40, 0.5 % deoxycholate) with proteinase and phosphatase inhibitors (Roche Diagnostics GmbH).

### Western Blot analysis

Protein concentration of cell lysates was determined using the Pierce™ BCA protein assay kit according to manufacturer’s instructions (ThermoFisher Scientific). Samples were subsequently subjected to SDS-PAGE and blotted on a nitrocellulose membrane. The following antibodies were used in 5 % low-fat milk in TBS-Tween buffer: Puromycin (ms, Millipore, 12D10, 1:10.000), β-actin (ms, Sigma, AC-74, 1:25.000). After incubation with HRP-conjugated secondary antibody (Invitrogen, 1:5000), the blot was developed using ECL solution (Merck Millipore). Films were used for detection (Amersham Biosciences).

### RNA isolation and qRT-PCR

Matrigel droplets were degraded in 0.5 % trypsin, 0.5 mM EDTA in PBS for 8 min at 37 °C. Trypsin was neutralized using cold KGM. After centrifugation for 5 min at 2000 rpm cells were washed with PBS. The pellet was resuspended in QIAzol (Qiagen) and frozen in liquid nitrogen. Sorted and freshly isolated keratinocytes were collected in QIAzol and frozen in liquid nitrogen. Samples were subjected to three freeze/thaw cycles (liquid nitrogen/ 37 °C water bath). Half of the total volume of QIAzol was added per sample. After incubation for 5 min at RT, 200 µl chloroform were added per 1 ml QIAzol. The samples were vortexed, incubated for 2 min at RT, and centrifuged at 10.000 rpm and 4 °C for 15 min. The aqueous phase was mixed with an equal volume of 70 % ethanol and transferred to a Rneasy Mini spin column (Qiagen). Total RNA was isolated with the Rneasy Mini Kit (Qiagen) according to manufacturer’s instructions. cDNA was generated using the iScript cDNA Synthesis Kit (Bio-Rad Laboratories Inc.). qRT-PCR was performed using Power SYBR Green master mix (Applied Biosystems) on a ViiA 7 Real-Time PCR System (Applied Biosystems). Primer sequences are listed below.

**Table 1:**
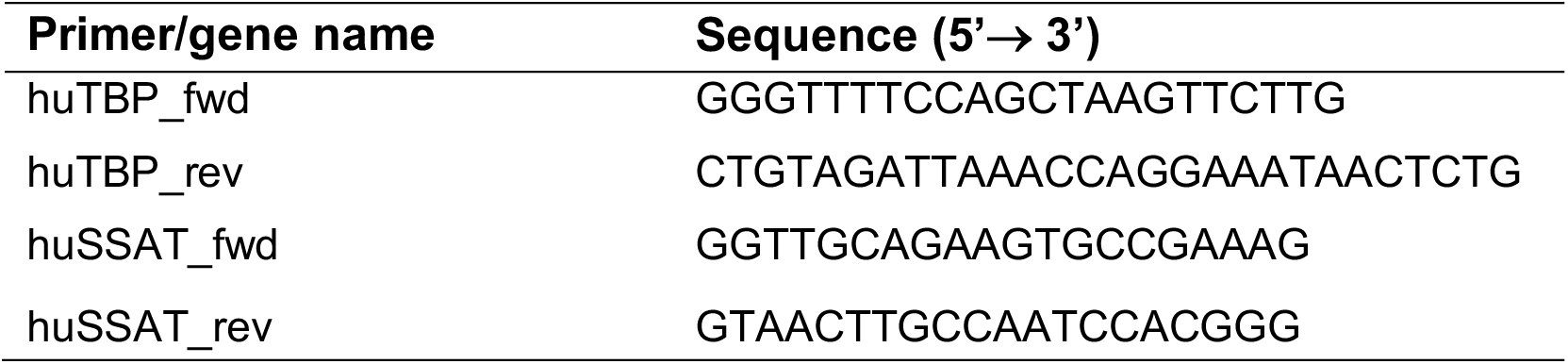
Primer sequences for qRT-PCR.

### Preparation of 3’ RNA-sequencing libraries

Sorted cells were collected in QIAzol and frozen in liquid nitrogen. Total RNA was isolated as described above. 50 ng RNA per sample were used for cDNA synthesis with Maxima H Minus reverse transcriptase (ThermoFisher Scientific). During reverse transcription, unique barcodes including unique molecular identifiers (UMI) were attached to each sample. After cDNA synthesis, all samples were pooled and processed in one single tube. DNA was purified using AmpureXP beads (Beckman Coulter) and the eluted cDNA was subjected to Exonuclease I treatment (New England Biolabs). cDNA was PCR-amplified for 12 cycles and subsequently purified. After purification, cDNA was tagmented in 5 technical replicates of 1 ng cDNA each using the Nextera XT Kit (Illumina), according to manufacturer’s instructions. The final library was purified and concentration and size were validated by Qubit and High Sensitivity TapeStation D1000 analyses. A detailed protocol for library preparation is available on request. Sequencing was carried out on an Illumina NovaSeq system.

### Bioinformatic analysis of 3’ RNA-sequencing data

The raw reads were demultiplexed and the UMI-tag was added to each read name using FLEXBAR (Dodt et al., 2012). The second end of the read-pairs were mapped using Kallisto creating pseudo-alignments to the mouse reference genome mm10 (Bray et al., 2016). UMI-tags were assigned using BEDTools intersect and bash re-formatting (Quinlan & Hall, 2010). An R script was used to count the unique UMI-tags per gene. Differential gene expression was calculated using the R package Deseq2 (Love et al., 2014). Only genes with an average of more or equal to 5 reads over all samples were used for the analysis. Only genes with an adjusted p-value ≤ 0.05 (α6+/CD34- vs. α6+/CD34+; untreated or treated) are shown in the heatmap. Venn diagrams were generated using the R package VennDiagram. Functional enrichment analysis was performed using the Metascape tool [http://metascape.org] (Zhou et al., 2019). Only genes with a p-value < 0.05 and a log2FC > ± 0.5 (α6+/CD34- vs. α6+/CD34+; untreated or treated) were defined as differentially expressed and included in the functional enrichment analysis.

### Polyamine extraction from cells

Matrigel droplets were degraded in 0.5 % trypsin, 0.5 mM EDTA in PBS for 8 min at 37 °C. Trypsin was neutralized using cold KGM. After centrifugation for 5 min at 2000 rpm cells were washed with PBS. Sorted cells were centrifuged and washed with PBS. The pellet was flash frozen in liquid nitrogen. The cells were lysed in ddH2O by freeze/thaw cycles and the protein concentration was determined using the Pierce™ BCA protein assay kit (ThermoFisher Scientific). Polyamines were extracted by Bligh and Dyer extraction (Noga et al., 2012, Pieragostino et al., 2015). In brief, a methanol:chloroform mixture 2:1 (v/v) was added to a volume of cell lysate, which corresponded to 100 µg of protein and incubated 1 h at 4 °C. Samples were centrifuged for 5 min at 3000 rcf at 4 °C and the liquid phase was transferred and dried in a SpeedVac Vacuum Concentrator (Genevac). Samples were reconstituted with 15 µL aqueous acetonitrile 2:3 (v/v) and 10 µl were injected into the LC-MS system.

### Targeted analysis of polyamines by LC-MS

The identification of polyamines was performed on a triple stage quadrupole (TSQ Altis, ThermoFisher Scientific GmbH) coupled with a binary pump system (Vanquish, ThermoFisher Scientific GmbH). Polyamine species were separated using a reverse column (Xselect column, 2.1×100mm, 2,5 µm) using solvent A (water with 0.1 % formic acid) and B (acetonitrile with 0.1 % formic acid) as previously reported (Hakkinen et al., 2008, Hakkinen et al., 2013, Sanchez-Lopez et al., 2009).

The gradient started from 0.1 % eluent B and ramped to 0.3 % eluent B in 0.5 min. It ramped further to 0.5 % eluent B in 0.5 min and to 1 % B in the next 0.5 min. The gradient increased to 2 % eluent B in the next minute, then it ramped to 5 % eluent B in 1 min. In the following 3 min it went to 95 % eluent B and stayed constant for 3 min. Afterwards the gradient decreased to 0.1 % eluent B in 2 min and stayed constant for 3 min adding up to a total time of 15 min. The column was heated to 30 °C using a flow rate of 100 µl/min. The LC system was flushed in between runs with isopropanol:water 75:25 (v/v) with 0.1 % FA.

Polyamines were detected using heated electrospray ionization (HESI) with the following parameters: 10 (a.u.) sheath gas, 5 (a.u.) auxiliary, 200 °C transfer ion capillary and 3 kV for spray voltage.

Relative quantification was performed using selected ion monitoring chromatogram mode (SIM-Q1) in positive ion mode using a scan rate of 1000 Da/sec, Q1 resolution was set to 0.7 *m/z* and calibrated RF lenses were used. The following ions were monitored: arginine → 175.2, ornithine → 133.17, putrescine → 89.17, spermidine → 146.23, spermine → 203.32, N1-acetylspermidine → 188.25, N1-acetylspermine → 245.34, N1,N11-diethylnorspermine → 245.39. The relative response for each polyamine was calculated using Spermidine-(butyl-d8) as internal standard.

**Table 2:**
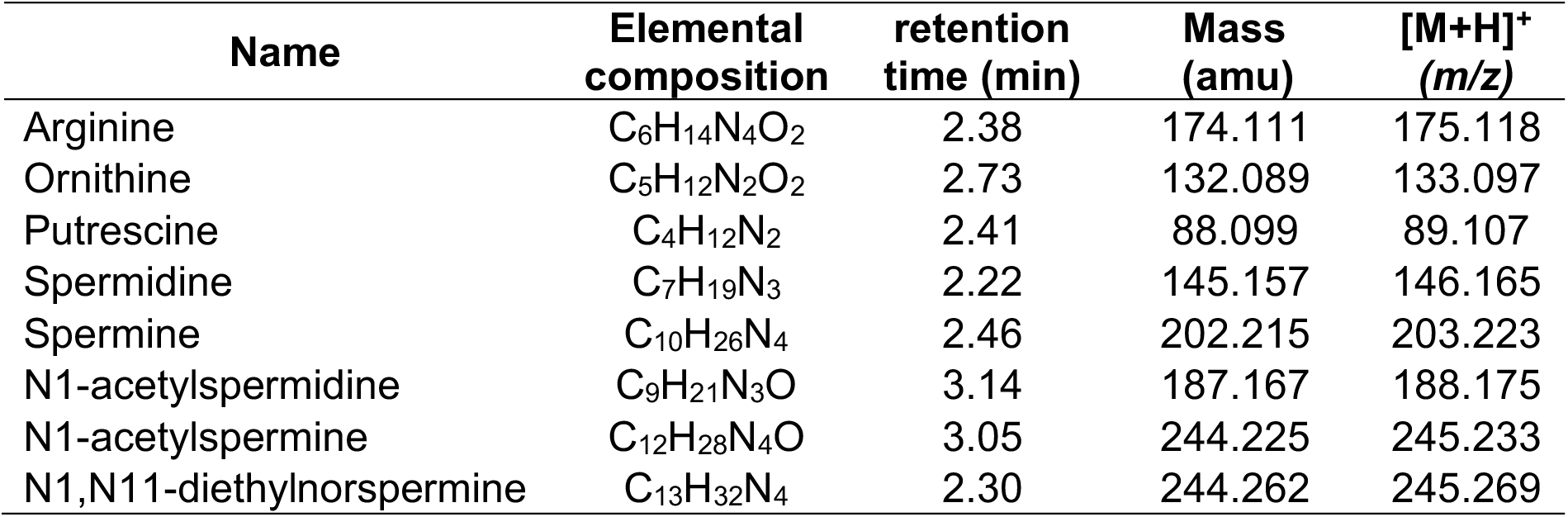
Detailed description of the detected metabolites.

| Name | Elemental composition | retention time (min) | Mass (amu) | [M+H]<sup>+</sup> (m/z) |
| --- | --- | --- | --- | --- |
| Arginine | C<sub>6</sub>H<sub>14</sub>N<sub>4</sub>O<sub>2</sub> | 2.38 | 174.111 | 175.118 |
| Ornithine | C<sub>5</sub>H<sub>12</sub>N<sub>2</sub>O<sub>2</sub> | 2.73 | 132.089 | 133.097 |
| Putrescine | C<sub>4</sub>H<sub>12</sub>N<sub>2</sub> | 2.41 | 88.099 | 89.107 |
| Spermidine | C<sub>7</sub>H<sub>19</sub>N<sub>3</sub> | 2.22 | 145.157 | 146.165 |
| Spermine | C<sub>10</sub>H<sub>26</sub>N<sub>4</sub> | 2.46 | 202.215 | 203.223 |
| N1-acetylspermidine | C<sub>9</sub>H<sub>21</sub>N<sub>3</sub>O | 3.14 | 187.167 | 188.175 |
| N1-acetylspermine | C<sub>12</sub>H<sub>28</sub>N<sub>4</sub>O | 3.05 | 244.225 | 245.233 |
| N1,N11-diethylnorspermine | C<sub>13</sub>H<sub>32</sub>N<sub>4</sub> | 2.30 | 244.262 | 245.269 |

### Data Availability

The raw RNA-sequencing data in this publication are available upon request from the authors.

## Acknowledgements

We thank all M.S.D. laboratory members for lively and helpful discussions. We thank F.A.M.C. Mayr for critical reading of the manuscript. We thank K. Folz-Donahue and L. Schumacher from the FACS and imaging core facility, F. Metge, A. Iqbal and J. Boucas from the Bioinformatics core facility and the Comparative Biology Facility at the Max Planck Institute for Biology of Ageing. We thank Patrick Wollek for expert mouse work. This study was supported by the Cologne Graduate School for Ageing Research (to K.A.), by the Deutsche Forschungsgemeinschaft (DFG, German Research Foundation) – Projektnummer 73111208 – SFB 829 (to S.A.W. and M.S.D.), by the Jane and Aatos Erkko Foundation (to S.A.W.), by the ERC-StG 640254 and ERC-PoC 768524 (to M.S.D.), and by the Max Planck Society (to Ad.An., P.T., S.A.W., and M.S.D.).

## Conflict of Interest

The authors declare no potential competing interests.

## Authorship

K.A., C.S.K., An.An., A.P., C.A.C.-M., P.T., S.A.W. and M.S.D designed the research. Ad.An. gave feedback throughout the project. K.A. (original draft), C.S.K., A.P., P.T., S.A.W. and M.S.D (review & editing) wrote the manuscript. An.An. and C.L. performed LC-MS analysis. A.P. prepared the library for 3’ RNA-sequencing. K.A. performed all other experiments. Funding acquisition: Ad.An., P.T., S.A.W. and M.S.D.

## Supplemental Figures

**Figure S1:**
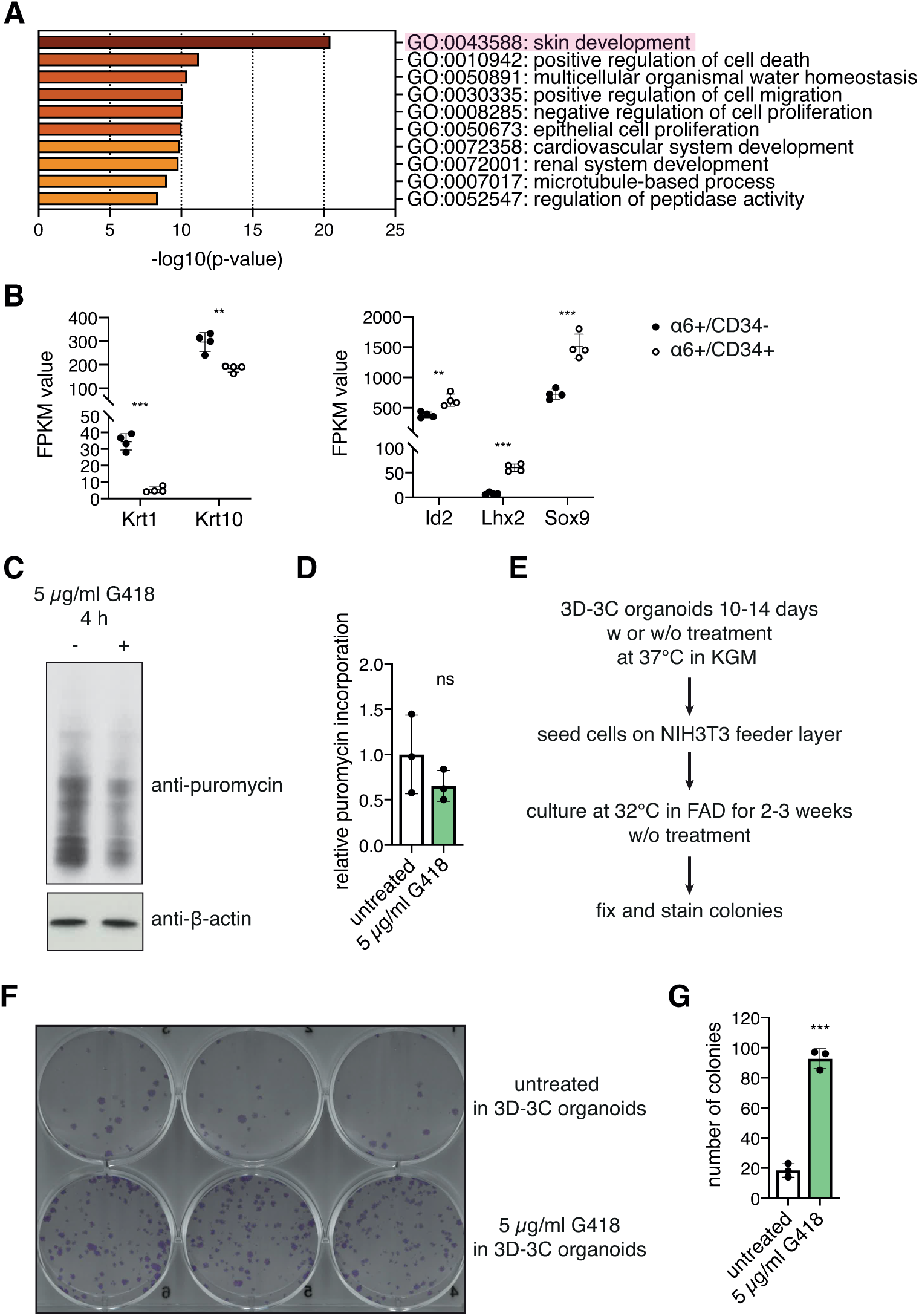
RNA-Sequencing confirms cell identity, which can be influenced by G418 treatment in the 3D-3C organoids. (A) GO term analysis of differentially expressed genes (p-value < 0.05, log2FC > ±0.5) from RNA-Seq experiment comparing α6+/CD34- progenitor cells with α6+/CD34+ stem cells (biological process, metascape.org). (B) FPKM values of differentiation (K1, K10) and stem cell marker genes (Id2, Lhx2, Sox9). Mean ± SD (n=4). *** p<0.001; ** p<0.01 (t test). (C) Representative Western blot analysis after puromycin incorporation in 3D-3C cultured cells. G418 treatment was performed for the last 4 h of the culture. (D) Quantification of the Western blot in (C). Mean ± SD (n=3). ns: not significant (t test). (E) Schematic representation of the workflow for colony formation assay after 3D-3C organoid culture. (F) Representative image of tissue culture plate after colony formation assay using cells with or without G418 treatment in 3D-3C organoids (n=3). (G) Quantification of colony number in (F). Mean ± SD (n=3). *** p<0.001 (t test).

**Figure S2:**
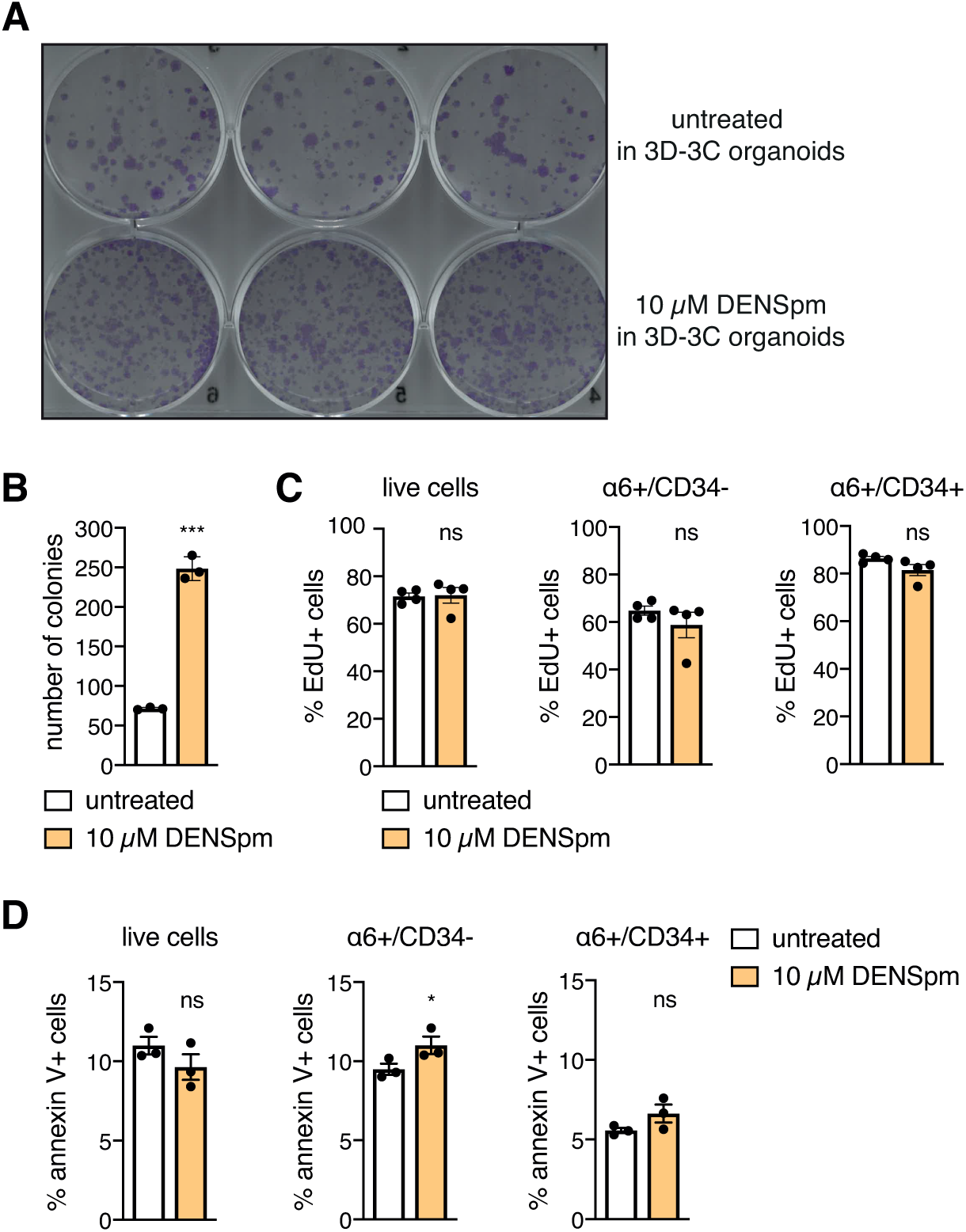
DENSpm treatment increases stem cell potency without major effects on proliferation or apoptosis. (A) Representative image of tissue culture plate after colony formation assay using cells with or without DENSpm treatment in 3D-3C organoids (n=2). (B) Quantification of colony number in (A). Mean ± SD (n=3). *** p<0.001 (t test). (C) Ratio of EdU+ cells with or without DENSpm treatment. EdU was incorporated for 24 h. Mean ± SEM (n=4). ns: not significant (t test). (D) Ratio of annexin V+ cells with or without DENSpm treatment. Mean ± SEM (n=4). * p<0.05; ns: not significant (t test).

**Figure S3:**
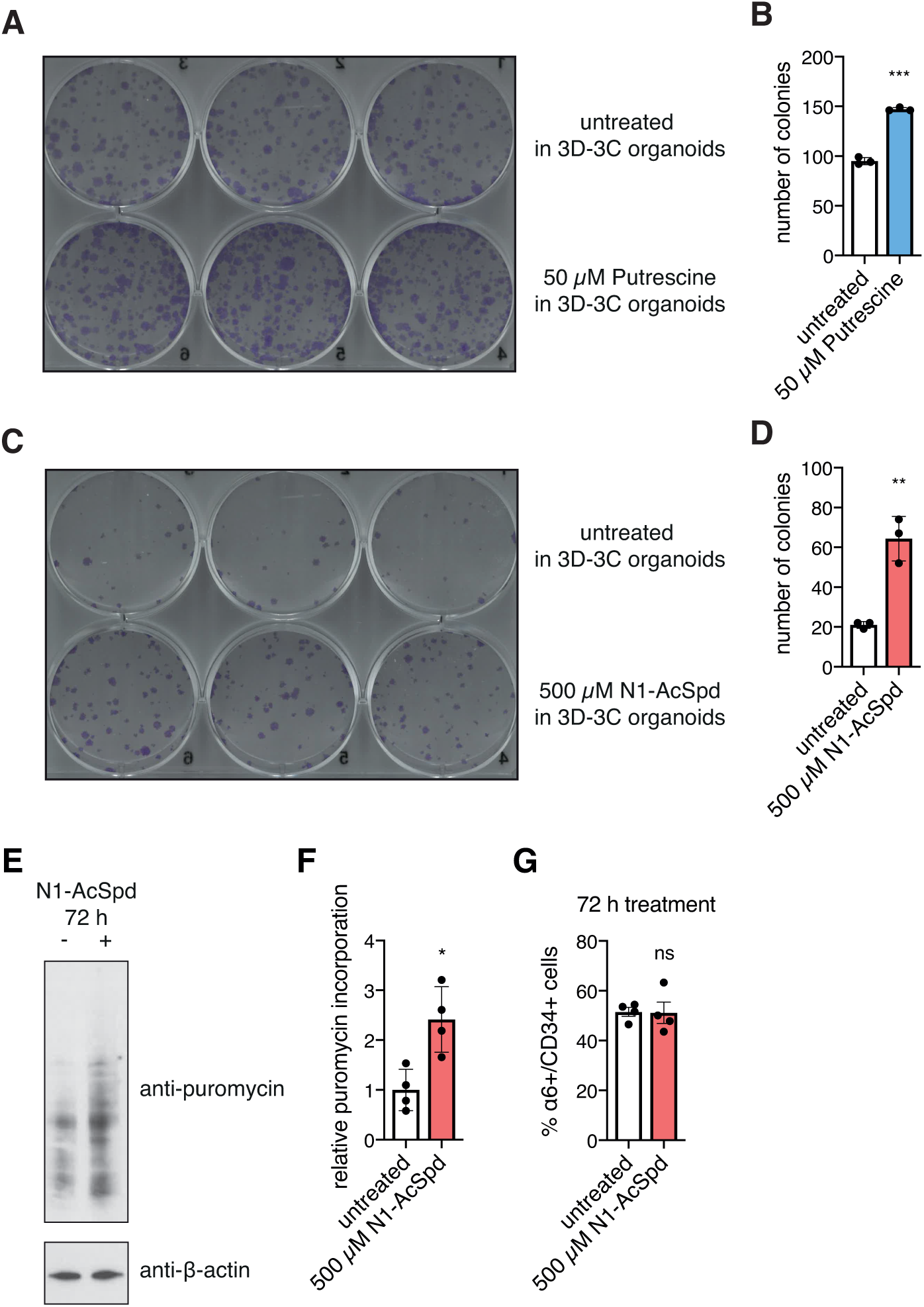
N1-acetylspermidine treatment increases stem cell potency without reducing translation. (A) Representative image of tissue culture plate after colony formation assay using cells with or without putrescine treatment in 3D-3C organoids (n=2). (B) Quantification of colony number of plate in (A). Mean ± SD (n=3). (C) Representative image of tissue culture plate after colony formation assay using cells with or without N1-AcSpd treatment in 3D-3C organoids (n=3). (D) Quantification of colony number of plate in (C). Mean ± SD (n=3). (E) Representative Western blot analysis after puromycin incorporation in 3D-3C cultured cells. N1-AcSpd treatment (500 µM) was performed for the last 72 h of the culture. (F) Quantification of the Western blot in (E). Mean ± SD (n=4). (G) Ratio of α6+/CD34+ cells after two weeks of 3D-3C culture with or without N1-AcSpd treatment for the last 72 h. Mean ± SEM (n=4). (A-G) Statistical significance was calculated by t test. *** p<0.001, ** p<0.01, p<0.05, ns: not significant.

**Figure S4:**
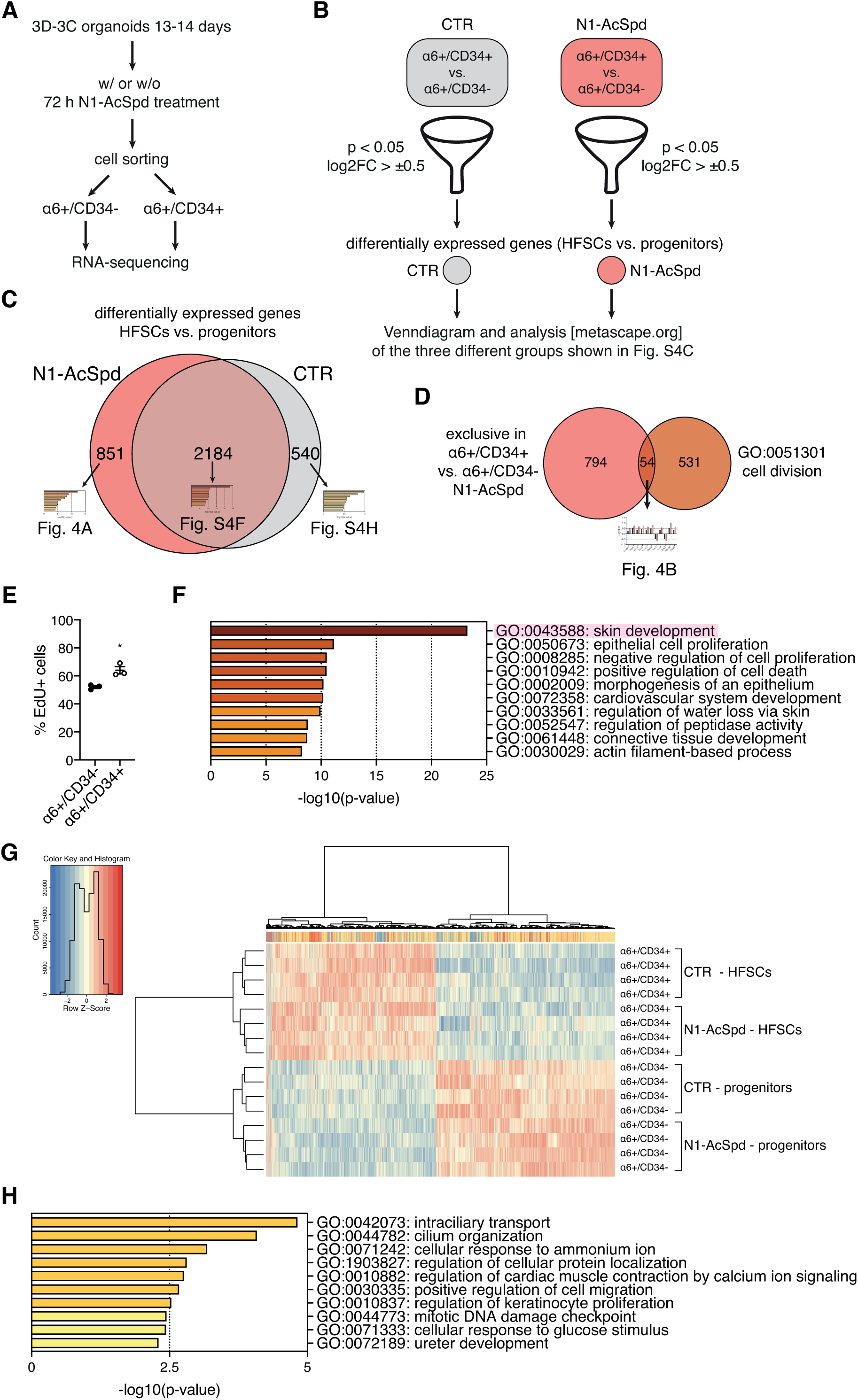
Schematic representation of the workflow of RNA sequencing analysis, which revealed maintained cell identity upon N1-acetylspermidine treatment. (A) Schematic representation of the workflow for RNA-seq sample collection. (B) Schematic representation of the bioinformatic workflow. α6+/CD34- cells and α6+/CD34+ cells were compared for untreated and treated conditions. Genes were filtered (p-value < 0.05, log2FC > ±0.5) and the resulting lists were used for further analysis. Control (CTR) is shown in gray, N1-AcSpd is depicted in red. (C) Venn diagram of the two groups from (B). CTR is shown in gray, N1-AcSpd is depicted in red. (D) Venn diagram of differentially expressed genes upon N1-AcSpd treatment (red) and genes covered by GO term cell division (brown). (E) Ratio of EdU+ cells in α6+/CD34- progenitors and α6+/CD34+ HFSCs. Mean ± SEM (n=4). * p<0.05 (t test). (F) GO term analysis of differentially expressed genes between cell types (p-value < 0.05, log2FC > ±0.5) from RNA-seq experiment common in untreated and treated cells (overlap shown in (C), biological process, metascape.org). (G) Heat map showing differentially expressed genes with a p-value ≤ 0.05 (α6+/CD34- vs. α6+/CD34+; treated or untreated). (H) GO term analysis of differentially expressed genes between cell types (p-value < 0.05, log2FC > ±0.5) from RNA-seq experiment only in untreated cells (gray in (C), biological process, metascape.org).

## Notes

### Competing Interest Statement

The authors have declared no competing interest.

## References

Bar-Nun S, Shneyour Y, Beckmann JS (1983) G-418, an elongation inhibitor of 80 S ribosomes. Biochim Biophys Acta 741: 123–7

Blanco S, Bandiera R, Popis M, Hussain S, Lombard P, Aleksic J, Sajini A, Tanna H, Cortes-Garrido R, Gkatza N, Dietmann S, Frye M (2016) Stem cell function and stress response are controlled by protein synthesis. Nature 534: 335–40

Blanco S, Kurowski A, Nichols J, Watt FM, Benitah SA, Frye M (2011) The RNA-methyltransferase Misu (NSun2) poises epidermal stem cells to differentiate. PLoS Genet 7: e1002403

Blanpain C, Fuchs E (2009) Epidermal homeostasis: a balancing act of stem cells in the skin. Nat Rev Mol Cell Biol 10: 207–17

Bray NL, Pimentel H, Melsted P, Pachter L (2016) Erratum: Near-optimal probabilistic RNA-seq quantification. Nat Biotechnol 34: 888

Calder A, Roth-Albin I, Bhatia S, Pilquil C, Lee JH, Bhatia M, Levadoux-Martin M, McNicol J, Russell J, Collins T, Draper JS (2013) Lengthened G1 phase indicates differentiation status in human embryonic stem cells. Stem Cells Dev 22: 279–95

Canellakis ZN, Marsh LL, Bondy PK (1989) Polyamines and their derivatives as modulators in growth and differentiation. Yale J Biol Med 62: 481–91

Chacon-Martinez CA, Klose M, Niemann C, Glauche I, Wickstrom SA (2017) Hair follicle stem cell cultures reveal self-organizing plasticity of stem cells and their progeny. EMBO J 36: 151–164

Clegg CH, Linkhart TA, Olwin BB, Hauschka SD (1987) Growth factor control of skeletal muscle differentiation: commitment to terminal differentiation occurs in G1 phase and is repressed by fibroblast growth factor. J Cell Biol 105: 949–56

Cohen SS, Lichtenstein J (1960) Polyamines and ribosome structure. J Biol Chem 235: 2112–6

Coleman CS, Huang H, Pegg AE (1995) Role of the carboxyl terminal MATEE sequence of spermidine/spermine N1-acetyltransferase in the activity and stabilization by the polyamine analog N1,N12-bis(ethyl)spermine. Biochemistry 34: 13423–30

Dever TE, Ivanov IP (2018) Roles of polyamines in translation. J Biol Chem 293: 18719–18729

Dodt M, Roehr JT, Ahmed R, Dieterich C (2012) FLEXBAR-Flexible Barcode and Adapter Processing for Next-Generation Sequencing Platforms. Biology (Basel) 1: 895–905

Fogel-Petrovic M, Vujcic S, Brown PJ, Haddox MK, Porter CW (1996) Effects of polyamines, polyamine analogs, and inhibitors of protein synthesis on spermidine-spermine N1-acetyltransferase gene expression. Biochemistry 35: 14436–44

Fredlund JO, Johansson MC, Dahlberg E, Oredsson SM (1995) Ornithine decarboxylase and S-adenosylmethionine decarboxylase expression during the cell cycle of Chinese hamster ovary cells. Exp Cell Res 216: 86–92

Fuchs E, Merrill BJ, Jamora C, DasGupta R (2001) At the roots of a never-ending cycle. Dev Cell 1: 13–25

Guo S, Zi X, Schulz VP, Cheng J, Zhong M, Koochaki SH, Megyola CM, Pan X, Heydari K, Weissman SM, Gallagher PG, Krause DS, Fan R, Lu J (2014) Nonstochastic reprogramming from a privileged somatic cell state. Cell 156: 649–62

Gutierrez E, Shin BS, Woolstenhulme CJ, Kim JR, Saini P, Buskirk AR, Dever TE (2013) eIF5A promotes translation of polyproline motifs. Mol Cell 51: 35–45

Hakkinen MR, Keinanen TA, Vepsalainen J, Khomutov AR, Alhonen L, Janne J, Auriola S (2008) Quantitative determination of underivatized polyamines by using isotope dilution RP-LC-ESI-MS/MS. Journal of pharmaceutical and biomedical analysis 48: 414–21

Hakkinen MR, Roine A, Auriola S, Tuokko A, Veskimae E, Keinanen TA, Lehtimaki T, Oksala N, Vepsalainen J (2013) Analysis of free, mono- and diacetylated polyamines from human urine by LC-MS/MS. Journal of chromatography B, Analytical technologies in the biomedical and life sciences 941: 81–9

Hershey JWB, Sonenberg N, Mathews MB (2019) Principles of Translational Control. Cold Spring Harb Perspect Biol 11

Hsu YC, Li L, Fuchs E (2014) Emerging interactions between skin stem cells and their niches. Nat Med 20: 847–56

Hsu YC, Pasolli HA, Fuchs E (2011) Dynamics between stem cells, niche, and progeny in the hair follicle. Cell 144: 92–105

Ingolia NT, Lareau LF, Weissman JS (2011) Ribosome profiling of mouse embryonic stem cells reveals the complexity and dynamics of mammalian proteomes. Cell 147: 789–802

James C, Zhao TY, Rahim A, Saxena P, Muthalif NA, Uemura T, Tsuneyoshi N, Ong S, Igarashi K, Lim CY, Dunn NR, Vardy LA (2018) MINDY1 Is a Downstream Target of the Polyamines and Promotes Embryonic Stem Cell Self-Renewal. Stem Cells 36: 1170–1178

Jensen UB, Yan X, Triel C, Woo SH, Christensen R, Owens DM (2008) A distinct population of clonogenic and multipotent murine follicular keratinocytes residing in the upper isthmus. J Cell Sci 121: 609–17

Kakegawa T, Guo Y, Chiba Y, Miyazaki T, Nakamura M, Hirose S, Canellakis ZN, Igarashi K (1991) Effect of acetylpolyamines on in vitro protein synthesis and on the growth of a polyamine-requiring mutant of Escherichia coli. J Biochem 109: 627–31

Kim CS, Ding X, Allmeroth K, Kolenc OI, L’Hoest N, Chacon-Martinez CA, Edlich-Muth C, Giavalisco P, Quinn KP, Denzel MS, Eming SA, Wickstrom SA (2019) An mTORC2-dependent switch to glutamine metabolism controls stem cell fate reversibility and long-term maintenance in the hair follicle. Manuscript submitted for publication

Kristensen AR, Gsponer J, Foster LJ (2013) Protein synthesis rate is the predominant regulator of protein expression during differentiation. Mol Syst Biol 9: 689

Landau G, Bercovich Z, Park MH, Kahana C (2010) The role of polyamines in supporting growth of mammalian cells is mediated through their requirement for translation initiation and elongation. J Biol Chem 285: 12474–81

Li A, Simmons PJ, Kaur P (1998) Identification and isolation of candidate human keratinocyte stem cells based on cell surface phenotype. Proc Natl Acad Sci U S A 95: 3902–7

Liu L, Michowski W, Inuzuka H, Shimizu K, Nihira NT, Chick JM, Li N, Geng Y, Meng AY, Ordureau A, Kolodziejczyk A, Ligon KL, Bronson RT, Polyak K, Harper JW, Gygi SP, Wei W, Sicinski P (2017) G1 cyclins link proliferation, pluripotency and differentiation of embryonic stem cells. Nat Cell Biol 19: 177–188

Love MI, Huber W, Anders S (2014) Moderated estimation of fold change and dispersion for RNA-seq data with DESeq2. Genome Biol 15: 550

Lu R, Markowetz F, Unwin RD, Leek JT, Airoldi EM, MacArthur BD, Lachmann A, Rozov R, Ma’ayan A, Boyer LA, Troyanskaya OG, Whetton AD, Lemischka IR (2009) Systems-level dynamic analyses of fate change in murine embryonic stem cells. Nature 462: 358–62

Metcalf BW, Bey P, Danzin C, Jung MJ, Casara P, Vevert JP (1978) Catalytic irreversible inhibition of mammalian ornithine decarboxylase (E.C.4.1.1.17) by substrate and product analogs. Journal of the American Chemical Society 100: 2551–2553

Nancarrow MJ, Nesci A, Hynd PI, Powell BC (1999) Dynamic expression of ornithine decarboxylase in hair growth. Mech Dev 84: 161–4

Noga MJ, Dane A, Shi S, Attali A, van Aken H, Suidgeest E, Tuinstra T, Muilwijk B, Coulier L, Luider T, Reijmers TH, Vreeken RJ, Hankemeier T (2012) Metabolomics of cerebrospinal fluid reveals changes in the central nervous system metabolism in a rat model of multiple sclerosis. Metabolomics : Official journal of the Metabolomic Society 8: 253–263

Odenlund M, Holmqvist B, Baldetorp B, Hellstrand P, Nilsson BO (2009) Polyamine synthesis inhibition induces S phase cell cycle arrest in vascular smooth muscle cells. Amino Acids 36: 273–82

Park MH, Cooper HL, Folk JE (1981) Identification of hypusine, an unusual amino acid, in a protein from human lymphocytes and of spermidine as its biosynthetic precursor. Proc Natl Acad Sci U S A 78: 2869–73

Parry L, Balana Fouce R, Pegg AE (1995) Post-transcriptional regulation of the content of spermidine/spermine N1-acetyltransferase by N1N12-bis(ethyl)spermine. Biochem J 305 (Pt 2): 451–8

Pegg AE (2016) Functions of Polyamines in Mammals. J Biol Chem 291: 14904–12

Pieragostino D, D’Alessandro M, di Ioia M, Rossi C, Zucchelli M, Urbani A, Di Ilio C, Lugaresi A, Sacchetta P, Del Boccio P (2015) An integrated metabolomics approach for the research of new cerebrospinal fluid biomarkers of multiple sclerosis. Molecular bioSystems 11: 1563–72

Quinlan AR, Hall IM (2010) BEDTools: a flexible suite of utilities for comparing genomic features. Bioinformatics 26: 841–2

Ray RM, Zimmerman BJ, McCormack SA, Patel TB, Johnson LR (1999) Polyamine depletion arrests cell cycle and induces inhibitors p21(Waf1/Cip1), p27(Kip1), and p53 in IEC-6 cells. Am J Physiol 276: C684–91

Roux PP, Topisirovic I (2018) Signaling Pathways Involved in the Regulation of mRNA Translation. Mol Cell Biol 38

Ruiz S, Panopoulos AD, Herrerias A, Bissig KD, Lutz M, Berggren WT, Verma IM, Izpisua Belmonte JC (2011) A high proliferation rate is required for cell reprogramming and maintenance of human embryonic stem cell identity. Curr Biol 21: 45–52

Saini P, Eyler DE, Green R, Dever TE (2009) Hypusine-containing protein eIF5A promotes translation elongation. Nature 459: 118–21

Sampath P, Pritchard DK, Pabon L, Reinecke H, Schwartz SM, Morris DR, Murry CE (2008) A hierarchical network controls protein translation during murine embryonic stem cell self-renewal and differentiation. Cell Stem Cell 2: 448–60

Sanchez-Lopez J, Camanes G, Flors V, Vicent C, Pastor V, Vicedo B, Cerezo M, Garcia-Agustin P (2009) Underivatized polyamine analysis in plant samples by ion pair LC coupled with electrospray tandem mass spectrometry. Plant physiology and biochemistry : PPB 47: 592–8

Schmidt EK, Clavarino G, Ceppi M, Pierre P (2009) SUnSET, a nonradioactive method to monitor protein synthesis. Nat Methods 6: 275–7

Seiler N, Dezeure F (1990) Polyamine transport in mammalian cells. Int J Biochem 22: 211–8

Sela Y, Molotski N, Golan S, Itskovitz-Eldor J, Soen Y (2012) Human embryonic stem cells exhibit increased propensity to differentiate during the G1 phase prior to phosphorylation of retinoblastoma protein. Stem Cells 30: 1097–108

Signer RA, Magee JA, Salic A, Morrison SJ (2014) Haematopoietic stem cells require a highly regulated protein synthesis rate. Nature 509: 49–54

Soler AP, Gilliard G, Megosh LC, O’Brien TG (1996) Modulation of murine hair follicle function by alterations in ornithine decarboxylase activity. J Invest Dermatol 106: 1108–13

Sonnenberg A, Calafat J, Janssen H, Daams H, van der Raaij-Helmer LM, Falcioni R, Kennel SJ, Aplin JD, Baker J, Loizidou M, et al. (1991) Integrin alpha 6/beta 4 complex is located in hemidesmosomes, suggesting a major role in epidermal cell-basement membrane adhesion. J Cell Biol 113: 907–17

Sunkara PS, Ramakrishna S, Nishioka K, Rao PN (1981) The relationship between levels and rates of synthesis of polyamines during mammalian cell cycle. Life Sci 28: 1497–506

Tahmasebi S, Amiri M, Sonenberg N (2018a) Translational Control in Stem Cells. Front Genet 9: 709

Tahmasebi S, Khoutorsky A, Mathews MB, Sonenberg N (2018b) Translation deregulation in human disease. Nat Rev Mol Cell Biol 19: 791–807

Trempus CS, Morris RJ, Bortner CD, Cotsarelis G, Faircloth RS, Reece JM, Tennant RW (2003) Enrichment for living murine keratinocytes from the hair follicle bulge with the cell surface marker CD34. J Invest Dermatol 120: 501–11

Tsai YH, Lin KL, Huang YP, Hsu YC, Chen CH, Chen Y, Sie MH, Wang GJ, Lee MJ (2015) Suppression of ornithine decarboxylase promotes osteogenic differentiation of human bone marrow-derived mesenchymal stem cells. FEBS Lett 589: 2058–65

Uimari A, Keinanen TA, Karppinen A, Woster P, Uimari P, Janne J, Alhonen L (2009) Spermine analogue-regulated expression of spermidine/spermine N1-acetyltransferase and its effects on depletion of intracellular polyamine pools in mouse fetal fibroblasts. Biochem J 422: 101–9

Wallace HM, Fraser AV, Hughes A (2003) A perspective of polyamine metabolism. Biochem J 376: 1–14

Yamashita T, Nishimura K, Saiki R, Okudaira H, Tome M, Higashi K, Nakamura M, Terui Y, Fujiwara K, Kashiwagi K, Igarashi K (2013) Role of polyamines at the G1/S boundary and G2/M phase of the cell cycle. Int J Biochem Cell Biol 45: 1042–50

Zhang D, Zhao T, Ang HS, Chong P, Saiki R, Igarashi K, Yang H, Vardy LA (2012) AMD1 is essential for ESC self-renewal and is translationally down-regulated on differentiation to neural precursor cells. Genes Dev 26: 461–73

Zhao T, Goh KJ, Ng HH, Vardy LA (2012) A role for polyamine regulators in ESC self-renewal. Cell Cycle 11: 4517–23

Zhou Y, Zhou B, Pache L, Chang M, Khodabakhshi AH, Tanaseichuk O, Benner C, Chanda SK (2019) Metascape provides a biologist-oriented resource for the analysis of systems-level datasets. Nat Commun 10: 1523

Zismanov V, Chichkov V, Colangelo V, Jamet S, Wang S, Syme A, Koromilas AE, Crist C (2016) Phosphorylation of eIF2alpha Is a Translational Control Mechanism Regulating Muscle Stem Cell Quiescence and Self-Renewal. Cell Stem Cell 18: 79–90

